# The Clade Displacement Index: how to detect horizontal gene transfers in unrooted gene trees

**DOI:** 10.1101/2021.06.24.449756

**Authors:** Michał Aleksander Ciach

**Affiliations:** Faculty of Mathematics, Informatics and Mechanics, University of Warsaw, 2 Banacha St., 02-097, Warsaw, Poland

## Abstract

While most genes of any organism are inherited vertically - i.e. from its parent organisms - sometimes they can be exchanged between unrelated species in a process known as the horizontal gene transfer (HGT). Studies of HGT contribute to our knowledge about the mechanisms of evolution, including the emergence of new pathogens, and a great deal of effort has been put into different methods of finding transferred genes. The golden standard of HGT detection is the analysis of the incongruence between the gene and the species trees. Those methods typically require rooted trees, in which the direction of evolution is known. Gene trees are typically unrooted, and rooting them is yet another step in HGT analysis, prone to errors which may lead to wrong conclusions. A natural question arises: can HGTs be detected in gene trees without rooting them at all?

It turns out that, for a particular, yet broad, class of transfers, the answer to this question is: yes. It also turns out that the same methodology can be applied to complement the bootstrap support in assessing the stability of gene tree topology. In this article, we present the Clade Displacement Index, a measure of shift of a given clade’s location between two trees. We derive algorithms to compute it and give several examples of its applications to HGT detection and gene tree stability analysis. We finish by pointing out directions for further studies and an example that shows that not all HGTs are detectable without knowing the location of the root of the gene tree.

A Jupyter Notebook with the implementation and applications of CDI described in this paper is available at https://github.com/mciach/CDI

## 1 Introduction

In recent years, numerous methods of detecting horizontally transferred genes— i.e. genes acquired from distantly related organisms—have been developed and applied in evolutionary studies [1–3]. It is believed that methods based on phylogenetic trees offer the most reliable approach to this task [4–6]. Those methods detect genes that are evolutionarily more similar to each other than what would be expected from the evolutionary relationships of their host species, as depicted in Fig. 1.

**Figure 1:**
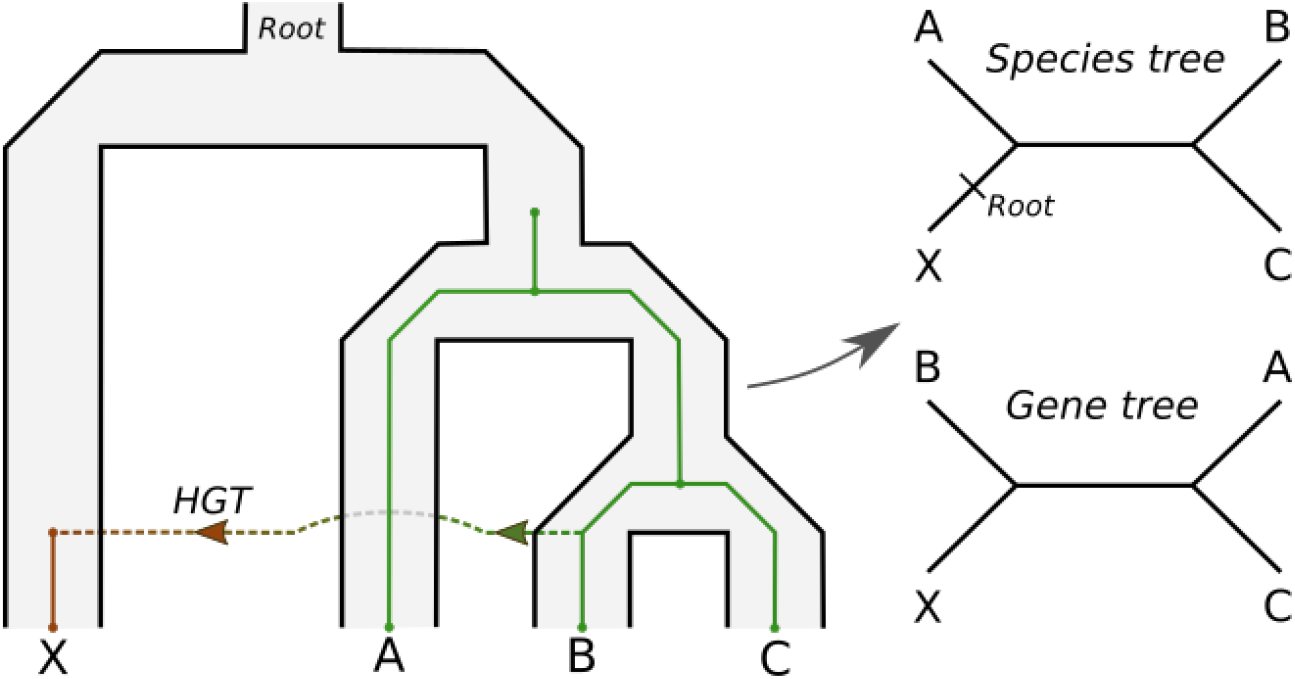
Horizontal gene transfer between distant species results in a different branching pattern in the gene and the corresponding species tree. The species X branches with the species A and the B/C clade in the species tree (after ignoring the root). However, the closest homolog to the gene that X received through HGT is the one from B, causing X to branch with B and A/C clade in the gene tree. This kind of incongruence between the gene and the species tree is a common argument for HGT.

Often, it is sufficient to analyze phylogenetic trees manually to infer an HGT event (see e.g. [7–10]). In case of more complex trees or larger data sets, algorithmic approaches are needed. An idea that has received a great deal of attention from both algorithmic and biological communities, with numerous mathematical models, implementations, and applications, is the tree reconciliation [11–17]. A common denominator of different approaches to tree reconciliation is to embed a gene tree, which represents an evolutionary history of a set of genes, into the corresponding species tree, which represents an evolutionary history of the species containing those genes. An example of such embedding is shown in Fig. 1.

The reconciliation may be done under a probabilistic model of gene tree evolution, where a maximum likelihood embedding is searched for, or in a parsimonious framework, where the costs of evolutionary events such as gene duplication or transfer are defined by the user, and a minimum-cost embedding is constructed. Although there are many approaches to tree reconciliation, most of the state-of-the-art algorithms that include the horizontal gene transfer in the set of possible evolutionary events share a common requirement that the gene tree needs to be rooted. Several studies have tackled the problem of unrooted reconciliations, but usually in a setting where gene duplications and losses are the only source of incongruence between trees [18, 19].

Unfortunately, trees obtained by the maximum likelihood (ML) method, widely considered as one of the most accurate, are typically unrooted, unless non-standard models of sequence evolution are used [20–25]. Such trees require rooting before further analysis. There is a number of rooting techniques with different scopes of applications and different drawbacks [26]. They can be roughly classified into two groups. The first group consists of methods based on branch lengths (which are proportional to the evolutionary distances between the nodes), such as the midpoint, minimal variance, and minimal ancestral deviation methods [27,28], which place the root in a way that balances out branch lengths on the two sides of the tree. However, such methods may fail when the molecular clock assumption (i.e. the assumption that the substitution rate is constant) is violated, which is the case for many gene trees.

The second group of rooting methods is based on known evolutionary relationships between some, or all, of the species. Two notable examples are the outgroup method, which uses a group of organisms distantly related to the organisms of interest and places the root between the two groups, and the minimum evolutionary cost method, which places the root in a way that minimizes said cost. The former is considered to be the most accurate as long as an appropriate outgroup is available, which is often not the case, especially in high-throughput analyses. The latter method is better suited for applications where the selection of an outgroup is either impossible or infeasible, but requires the user to specify the costs of evolutionary events. At this point, to our knowledge, there is a lack of systematic and comprehensive studies on how to properly estimate these costs and the impact of their misspecification, although some works start to appear [17, 29, 30]. Furthermore, this method often returns multiple root candidates, and is sensitive to errors in gene tree reconstruction [19, 31]. Overall, the correct placement of the root of a gene or a species tree is far from straightforward, sometimes to the point that detailed studies for single groups of organisms are needed [32, 33].

It turns out that when a gene is transferred between distantly related species, such that the recipient organism does not contain other homologs of the transferred gene, the resulting incongruence between the gene and the species tree may be sufficiently simple and clear that HGT can be inferred without rooting. An example of such transfer is depicted schematically in Fig. 1, and actual transfers of this kind were studied from a biological perspective e.g. in [7, 8]. In those cases, the recipient organism—or a group of organisms in case of an ancestral transfer—branches with different clades in the gene and the species tree. This is because its genes are closely related to the ones of the donor organism, causing the recipient clade to branch with the donor in the gene tree.

Based on the above observation, in this article we propose a method of detecting such events in unrooted gene trees. The method is based on comparing the location of a given clade in a gene tree and the corresponding species tree. If we detect that a clade of interest has a different location in both trees, we may suspect that the last common ancestor of this clade is an acceptor of a horizontally transferred gene.

Naturally, other evolutionary events, such as gene duplications, and populational effects, such as incomplete lineage sorting, may change the topology of the gene tree, sometimes to the point where it is hardly similar to the corresponding species tree. However, our main area of interest is genome screening for clear-cut cases of HGT, where the goal is to detect horizontally transferred genes in the genome of an organism of interest with a small false positive rate. Therefore, we focus on fairly simple evolutionary scenarios, with a single transfer event, and such that other phenomena have a relatively small impact on the topology of the gene tree.

We start with a simple example to illustrate the idea behind our approach. Then, we generalize our approach so that it can be applied to any pair of non-binary, non-uniquely labeled trees. We then show that this general approach can be simplified in the case of comparing gene and species trees, the case most common in phylogenetics. Interestingly, the time complexity of our method changes from linear in the simplest case, to exponential in the general case, to linear again in the case of gene and species trees. We finish the article by showing two applications of our method. First, we show the intended application to HGT detection, together with a control example in which no HGT occurs. Second, we show that our method can also be used to complement the bootstrap support when assessing the stability of the topology of a tree, a somewhat unexpected application.

## 2 Methods

### 2.1 Definitions

We start with the formal definitions of terms and symbols used throughout this article. A *binary unrooted tree* is a connected, undirected, acyclic graph such that the degree of each internal node equals 3. Since in this work we only consider unrooted trees, we will refer to them simply as trees. A tree is *non-binary* if any internal node has a degree larger than 3. Such nodes are referred to as *polytomous*. A *labeling* of leaves of a tree *T* is a surjective function from the set of all leaves *L*(*T*) onto a set of labels *ε*(*T*). We denote the number of *e*-labeled leaves of *T* as *T*(*e*). We say that the labeling is *unique* if *T*(*e*) = 1 for all *e* ∈ *ε*(*T*).

A *clade* is a group of nodes of a given tree that can be separated from all the remaining nodes by removing a single branch. The node adjacent to this branch that belongs to the clade is referred to as the clade’s *root* node. We say that a clade is *attached* to a node if its root is adjacent to this node. A clade is a *neighbour* of another clade if they are attached to the same node. For example, in Fig. 1, the neighbouring clades of the single-leaf clade X in the species tree are the leaf A and the subtree induced by leaves B and C. A clade is *induced* by a set of leaf labels if it contains all the leaves with those labels and no leaves with other labels. Note that a given set of leaf labels may not induce a clade. For example, in Fig. 1, labels A and C induce a subtree in the species tree, but not a clade.

### 2.2 The Simple Case

In order to give the reader an intuition behind our approach, we start with a simple case of two binary trees with a unique leaf labelling, *T*_1_ and *T*_2_, with identical sets of leaf labels *ε*(*T*_1_) = *ε*(*T*_2_). Consider two clades, 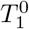 in *T*_1_ and 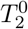 in *T*_2_, which locations are to be compared. We will refer to 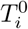 as the *reference* clades.

Throughout this article, we assume that the reference clades also have identical sets of leaf labels 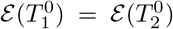. This is motivated by the intended application of our approach, where we compare the locations of clades induced by a given set of species in gene and species trees. In practice, this assumption may be relaxed by simply ignoring leaves with non-shared labels in both trees.

Since the trees *T_i_* are binary, each clade admits two neighbouring clades. Designate the neighbouring clades of 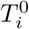 as 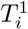 and 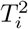. Note that, since we’ve assumed that 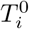’s have a common set of labels, we have 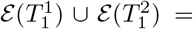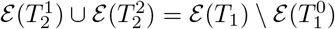.

Our notion of the difference of the location of reference clades in both trees, or the *Clade Displacement Index* (CDI), is based on comparing the leaf labels of their neighbouring clades. Namely, we consider the reference clades 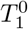 and 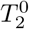 to be similarly located in *T*_1_ and *T*_2_ if their neighbouring clades induce a similar partition of the set of leaf labels 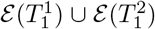. This approach is free from problems arising from possible differences in tree topology, which occur naturally due to e.g. errors in tree inference procedures, and allows to define the clade displacement rigorously.

Formally, we will measure the difference of location of 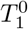 and 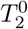 by matching the 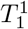 clade to one of the two 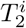 clades and the 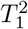 clade to the other, and measuring the difference in label composition of the matched subtrees. The matching is done in a way to minimize said difference. We can formulate this idea with the following equation, where *A*Δ*B* denotes the symmetrical difference of sets *A* and *B*:

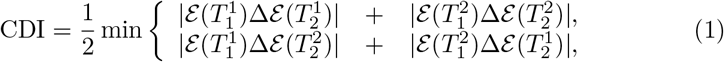

where the minimum is taken over the two rows.

It turns out that we do not need the two summands in the sums above, and it suffices to analyze the disagreement betweem 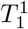 and one of the 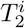 clades. One can easily show that 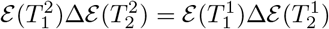, and therefore

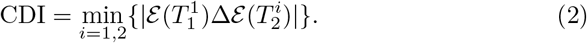

This equation serves as a basis for a simple linear time algorithm to obtain the Clade Displacement Index in the particular case presented in this subsection.

In its essence, CDI is a measure of similarity of splits—partitions of the set of leaf labels—induced by the neighbouring clades in both trees. In this way, it bears some resemblance to the well-known concept of the Robinson-Foulds distance [34] between two trees. This distance considers all the possible splits induced by edges of both trees, and is defined as the size of the symmetrical difference of the sets of splits. On the other hand, in CDI, in each tree we consider a single split, but compare the two splits in a more detailed manner. Somewhat similar approaches to split comparison have been used to generalize the Robinson-Foulds distance and make it less sensitive to minor changes in tree topology [35].

We will now rewrite Eq. (2) in a way that will serve as a basis for a generalization into trees with a non-unique labeling. With each 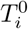 we associate two variables 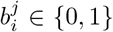, and we will match 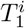 with 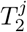 when 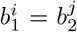. We require that, for each *i*, exactly one of 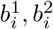 is equal to 1, and the other is equal to 0, to guarantee that the neighbouring clades of 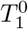 and 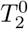 are matched one to one. For simplicity, we will refer to the clades with 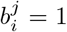 as the *left* clades, and to the ones with 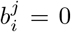 as the *right* clades. Using this notation, we can express CDI in the following way:

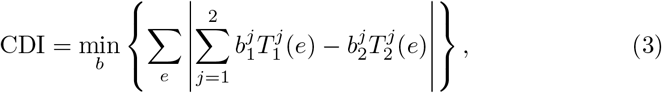

where the *b* variables need to satisfy the conditions described above, and the first sum goes over all labels 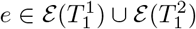.

Formulas (2) and (3) are equivalent, in the sense that they return the same value when applied to a given pair of trees. This can be shown as follows. Assume that the vectors 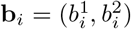 minimize the sum from Eq. (3). Observe that, for a vector 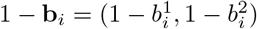, we have

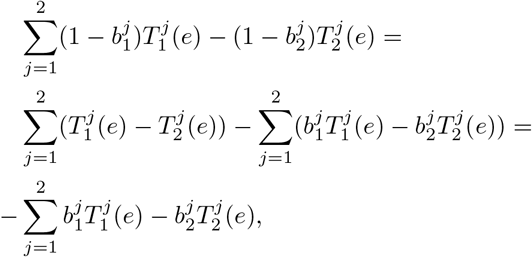

and, after taking the absolute value, we conclude that the vectors 1 − **b**_*i*_ and **b**_*i*_ yield equal values of CDI. The formulation of CDI in Eq. (3) is therefore invariant with respect to simultaneous swapping of left and right subtrees in both trees. In particular, we may fix the vector **b**_1_ arbitrarily as **b**_1_ = (1, 0), i.e. designate 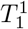 as the left subtree of *T*_1_. Equation (3) now simplifies to

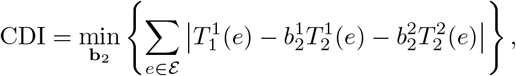

where exactly one of 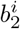 is equal to 1, and the other to 0. We therefore have

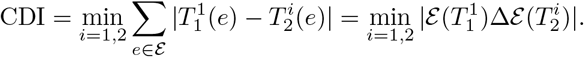

In the next sections, we will generalize Eq. (3) to handle non-binary, non-uniquely labeled trees. We will then reduce the general version again to get a more efficient algorithm for gene and species trees - where typically the former contains non-unique labels, while the latter is non-binary.

### 2.3 The General Case

In this section, we consider a general case of comparing the location of given clades in two non-binary trees with non-unique leaf labelling. In this case, comparing clade location is expressed as an integer programming problem.

Let *T*_1_, *T*_2_ be possibly non-binary trees with a possibly non-unique leaf labeling, with identical sets of leaf labels: *ε*(*T*_1_) = *ε*(*T*_2_). Let 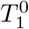 and 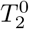 be the reference clades which locations are to be compared. Let 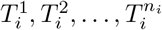 be the neighbouring clades of 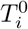. As before, we assume that 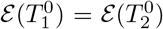, so that 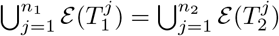.

We assume that a polytomous node represents a binary relationship with an unknown branching order. Consistently, for each *i* = 1, 2, we will merge the neighbouring clades 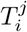 into two sets, referred to as the *left* and the *right* subtree. We will then compare the partitions of the set of labels induced by the merged neighbouring clades in both trees. The sets of labels obtained this way correspond to binarizations of the polytomous nodes to which 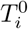’s are attached.

With each 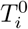 we associate a binary (column) vector 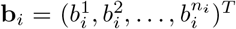 such that 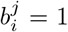 if 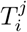 is put into the left subtree, and 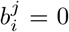 if 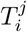 is put into the right one. In order to correspond to a proper binarization, the vector **b**_*i*_ needs to satisfy the condition 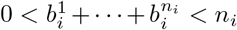, i.e. both the left and right subtree contains at least one of the neighbouring clades 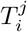.

In order to handle non-unique labeling, we define the *weight* of a label in a given subtree as the proportion of leaves with this label that fall into this subtree: 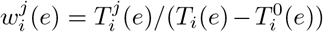. Note that 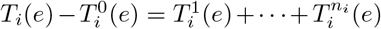 is the number of leaves with label *e* in the neighbouring clades. Putting the label weights in a column vector 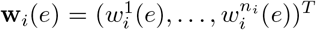 and multiplying by a binarization vector **b**_*i*_ yields a *left subtree weight* of a label,

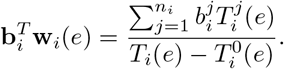

Combining the two ideas allows us to generalize Eq. (3) to the case of non-binary, non-uniquely labeled trees:

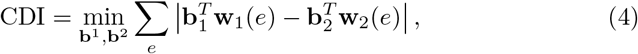

where the vectors **b**_*i*_ need to satisfy appropriate conditions for binarizations described earlier, and the first sum goes over all labels in 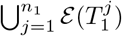. Note that the optimization problems (3) and (4) are equivalent in the case of uniquely labeled binary trees.

Problem (4) can be solved by expressing it as an instance of integer programming. First, introduce auxiliary variables 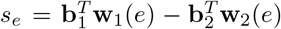 and express problem (4) as

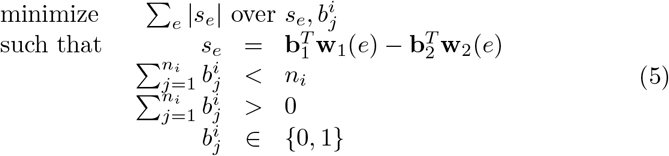

A standard approach to convert a problem like the one above into an instance of linear programming (see e.g. [36]) is to introduce new variables 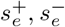 such that 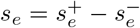, and minimize 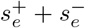 instead of |*s_e_*|:

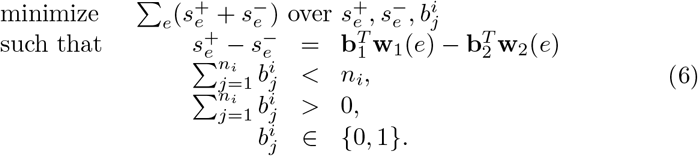

An optimization problem like the one above can be solved numerically by a number of linear program solvers. Note that, in general, integer programming is an NP-complete problem. At this point, we do not know if a polynomial time complexity algorithm for problem (4) exists. However, in the next section we present an efficient algorithm for the case of species tree-gene tree pairs.

### 2.4 The Case of Gene and Species Trees

A *species tree S* is a possibly non-binary, unrooted tree with a unique labelling. We refer to the labels of this tree as *species*. Note that our definition differs from the most common one, as the species trees are typically rooted; the reason for such definition is that we want to ignore the root location in *S*. A *gene tree G* corresponding to a species tree *S* is a binary tree with a possibly non-unique labeling of leaves such that *ε*(*G*) = *ε*(*S*).

Let 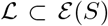 be a subset of species that induces clades *S*^0^ in *S* and *G*^0^ in *G*. Clades induced by 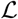 are our reference clades which locations are to be compared in both trees. As explained in the introduction, if such clade is displaced in the gene tree, we can suspect that the last common ancestor of this clade—correspoding to the clade’s root node in the species tree—obtained the gene through a horizontal gene transfer.

Since *G* is binary, *G*^0^ has two neighbouring clades, *G*^1^ and *G*^2^, with possibly non-disjoint sets of leaf labels. On the other hand, *S*^0^ may have multiple neighbouring clades *S*^1^, *S*^2^*, …, S^k^*, but their sets of leaf labels are pairwise disjoint due to unique labeling.

In the same way as for the simple case of binary, uniquely-labeled trees, one can show that CDI is symmetrical with respect to simultaneous swapping of left and right subtrees in both trees. We may therefore fix *G*^1^ as the left neighbour of *G*^0^ to obtain a fixed left tree score of labels in *G*, equal to 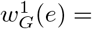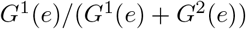 for *e* ∈ *ε*(*G*).

Consistently with the approach from the previous sections, we want to group the neighbouring clades in *S* into left and right subtrees in a way that maximizes the agreement with 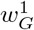. Consider a binarization vector **b** = (*b*^1^, *b*^2^*, …, b^k^*)^*T*^ such that *b^i^* = 1 if *S^i^* is put into the left subtree, and *b^i^* = 0 otherwise. Now, observe that each clade *S^i^* can be put into the left subtree independently of the others, in the sense that this operation changes the left tree weights for the labels from *S^i^* but not the other neighbouring clades. Formally, if *e* ∈ *ε*(*S^i^*), we have **b**^*T*^**w**_*S*_ (*e*) = *b^i^*, and therefore we can express CDI as

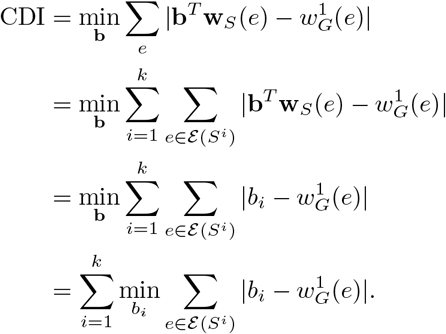

We can further simplify the equation above by noting that 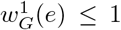, and therefore, if *b_i_* = 1, we have

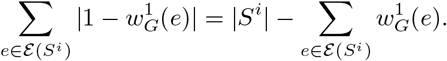

To get the value of CDI, we compute *k* partial scores 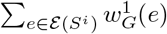 for *i* = 1, 2, …, *k*, compare them with 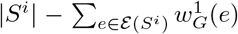, and sum up the lower ones. This algorithm has a linear time complexity in the number of leaves in *G* and *S* and the number of neighbouring clades in *S*.

## 3 Examples

### 3.1 Horizontal gene transfer in a parasitic plant

We illustrate the application of our approach on a previously described case of a horizontal gene transfer between a parasitic plant *Striga hermonthica* and one of its hosts, *Sorghum bicolor* [8]. We have accessed the predicted protein sequence of the putatively transferred gene from NCBI Protein database (GenBank accession BAJ06366.1), found its homologs in the protein NR database using the BlastP suite (https://blast.ncbi.nlm.nih.gov/Blast.cgi; sequences accessed and blasted on April 21, 2021) and selected 8 homologs from different species, reflecting the species composition from the original study. The selected species include the putative *Sorghum* donor and more distantly related *Triticum* and *Brachypodium* species. As a control example, we have downloaded the sequence of asparagine synthetase enzyme from *Striga hermonthica* (GenBank accession AAM94340.1) and selected 9 of its homologs found by BlastP.

For both data sets, have aligned the sequences using the MAFFT software (`--localpair --maxiterate=1000`), selected the conserved blocks using GBlocks (`-b5=h`), and constructed maximum likelihood trees with model selection using IqTree [22, 37, 38]. The BAJ06366.1 gene tree has been rooted by setting the *Triticum* and *Brachypodium* as an outgroup, and the AAM94340.1 tree has been rooted by midpoint method. Note that the trees are rooted only for the purposes of visualization and validation, as the root location is ignored by algorithms used in this study.

We have labeled the leaves of the gene trees with the names of their corresponding species names, downloaded the species trees from the NCBI Taxonomy (accessed 29.04.2021) and computed the branch displacement index of the *Striga* clades. The constructed trees, together with their corresponding species trees, are shown in Fig. 2.

**Figure 2:**
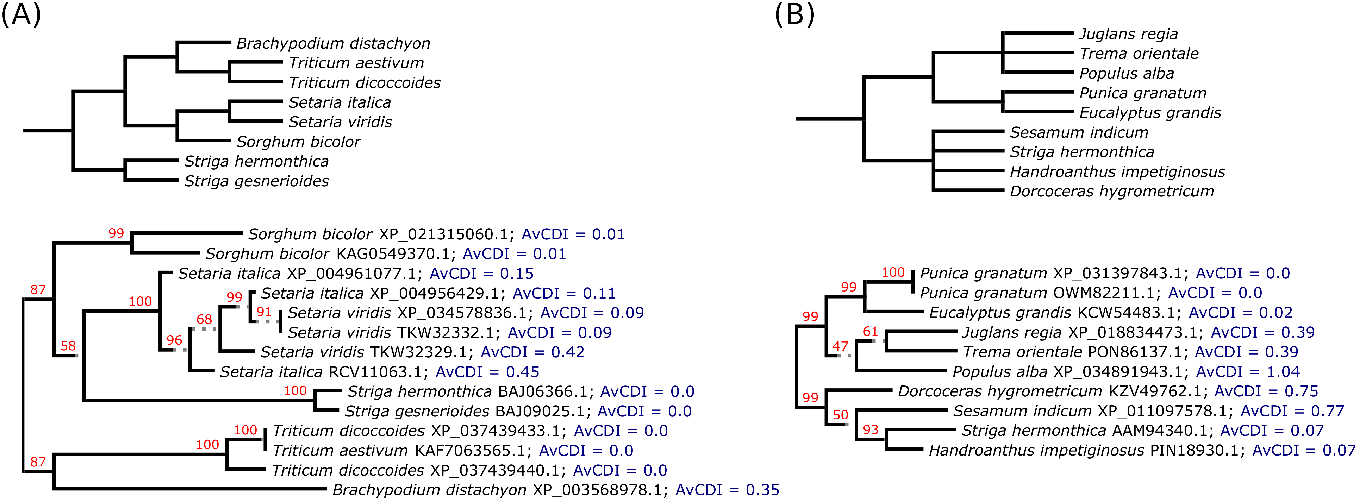
Example applications of the Clade Displacement Index. (A) The gene tree of BAJ06366.1 protein (bottom) and the corresponding species tree (up). The *Striga* species are an outgroup in the species tree, but branch with *Setaria* in the gene tree, indicating a horizontal gene transfer from the latter to the former clade. (B) The gene tree of AAM94340.1 protein (bottom) and the corresponding species tree (up). All clades have the same locations in both trees. AvCDI: Average Clade Displacement Index of a leaf in 100 bootstrap replicates. Numbers above branches indicate bootstrap supports.

In the first example, the *Striga* clade (consisting of two species) branches with *Sorghum* clade, supporting the HGT hypothesis. Accordingly, the displacement index of this clade is greater than zero (equal to 1), indicating a displacement of this clade with respect to the species tree. In the second example, the topologies of the gene and the species tree agree, in the sense that the polytomies of the species tree can be resolved in an order that matches the one of the gene tree. Accordingly, the displacement index of the *Striga* clade (consisting of one species) is zero.

The two examples presented above show that CDI can be used to detect a displacement of a clade of interest in a gene tree with respect to the corresponding species tree. Due to its efficient time complexity and no requirement to root the gene tree, it can be used to screen large datasets for putative HGT events.

### 3.2 Clade location consistency

Traditionally, bootstrapping is used to assess the reliability of the tree topology. The idea behind this technique can be summed up as randomly selecting columns from the alignment and inferring a phylogenetic tree from those columns. This procedure is repeated for a number of times. Then, for each branch of the reference tree, we inspect how often the split (i.e. the partition of leaf labels) induced by this edge occurs in the bootstrapped trees.

Although useful and used commonly, this technique has some major short-comings that make the interpretation of bootstrap supports difficult in some applications. Most importantly, the bootstrap support of an edge measures whether clades adjacent to this edge exist in bootstrapped trees, regardless of their location. Checking whether a given split is found in a tree is equivalent to checking whether this tree contains clades induced by the subsets of leaf labels corresponding to this split. How that can be misleading is shown in Fig. 3.

**Figure 3:**
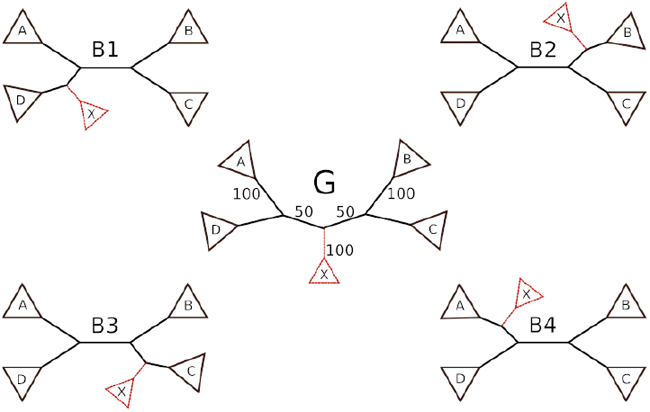
Bootstrap support values measure the consistency of clade existence, but not of its location. In a simplified scenario with a gene tree G and four bootstrap replicates from B1 to B4, the clade X exists in all of the replicates. Therefore, it has a full bootstrap support (100%), even though it is located differently in all the trees. The change in location needs to be inferred from a low bootstrap support (50%) of two of its adjacent branches. On the other hand, CDI allows to readily detect an inconsistency of clade location.

The branch displacement index offers an interesting complement to bootstrap when assessing the stability of leaf locations. While the bootstrap support of any leaf is always 100% (because a clade induced by a single label always exists in any bootstrapped tree), the branch displacement index shows whether it branches with similar clades, i.e. whether its location is stable in bootstrap replicates.

To demonstrate the application of this approach, we have computed 100 bootstrap replicates for both trees from the previous example. For each leaf of the gene tree, we have computed the mean and standard deviation of its displacement index (in this case referred to as the leaf displacement index) over the bootstrap replicates. Note that, in this case, we label the leaves of the gene tree and the bootstrapped trees with the protein accessions, not the species names, because we want to distinguish between leaves corresponding to different proteins of the same species. The species tree is not used in this analysis, because we’re only interested in the stability of the gene tree topology.

The results of this analysis are shown in Fig. 2. For the HGT example, the location of most leaves is stable, save for the RCV11063.1 protein of *Setaria italica* and TKW32329.1 of *Setaria viridis*. A closer inspection of the neighbouring clades shows that 68 replicates support the location from Fig. 2, while in 27 replicates TKW32329.1 and RCV11063.1 switch places. Those two scenarios constitute the vast majority of bootstrap replicates (95 out of 100).

The 100% bootstrap support of the *Striga* clade (i.e. the clade consisting of BAJ06366.1 and BAJ09025.1 proteins) means that this clade exists in all bootstrap replicates. Therefore, we can use the bootstrap replicates to check the stability of the location of the whole clade. It turns out that although the existence of this clade is well supported, its location is not. The bootstrapped displacement index of the *Striga* clade in the ML tree is zero in only 58 replicates, is equal to 2 in 23 replicates, and 4 in 10 replicates (with an overall average CDI equal to 1.05). Notably, in 23 replicates with CDI=2, the *Striga* clade branches with the *Triticum*/*Brachypodium* clade, in which case its location agrees with the one in the species tree and (without the knowledge of the root location of the gene tree) there is no support for the horizontal gene transfer.

In the second example, although the branching pattern of the ML gene tree agrees with the species tree, it is much less supported by the data in terms of bootstrap support and the average leaf displacement index. This is especially the case with *Populus alba*, with an average displacement index of 1.04. Accordingly, only 35 bootstrap replicates support its location on the ML tree. In 26 replicates, it branches with *Trema orientale* (a nearest neighbor interchange with CDI=1), in 17 replicates it branches with the *Eucalyptus*/*Punica* clade (with CDI=2), and in 21 replicates it branches with the whole *Eucalyptus*/*Punica*/*Juglans*/*Trema* clade (with CDI=2).

The two examples discussed in this section demonstrate that CDI can be used to assess the stability of the topology of gene trees. It complements and extends the capabilities of bootstrap support by allowing to inspect the stability of location of leaves (which always have 100% bootstrap support) and has a simpler interpretation, since it directly reflects the location of clades.

## 4 Discussion

We have presented a measure of similarity of the location of a given clade between two unrooted trees which share their leaf labels. Our goal when developing this approach was to be able to detect horizontal gene transfer events in a given set of species without the need to root gene trees. This requirement, common in tree reconciliation algorithms, is a major obstacle in high-throughput analyses. The current methods of rooting gene trees either require a manual approach (such as the outgroup method), are unreliable (such as the midpoint method), or require to specify a set of unknown parameters and may produce multiple root candidates (such as the minimum evolutionary cost approach). On the other hand, phylogenetic trees are the most informative structures that can be used to detect HGT, and methods based on phylogenies are considered to be more reliable than those based on sequence similarity or other sequence features. This is because the similarity of sequences may sometimes inaccurately reflect their evolutionary relationships, due to phenomena such as different substitution rates in different evolutionary lineages. Therefore, a method of detecting HGTs in unrooted gene trees is desirable.

Our approach is based on analyzing the label composition of clades neighboring with the clade of interest. In the case of unrooted binary trees, each clade has two other neighboring clades. We select one of the trees and arbitrarily designate one of the clades as the “left” one. Then, for each label, we calculate the proportion of leaves with this label that fall into the left clade. This yields a “left subtree score” for each label. In the second tree, we check which clade, when designated as the left one, will return a minimal sum of absolute differences of the left subtree scores over all labels. This sum is referred to as the Clade Displacement Index (CDI).

When the trees are non-binary, we partition the neighbouring clades into two sets to obtain a binarization of the polytomous node to which the clade of interest is attached. The binarization is done in a way that minimizes the displacement index. This approach is based on the assumption that polytomous nodes represent unknown order of evolutionary events, and the true evolutionary relationships can be represented by binary trees. In the general case of non-binary trees with multiply occurring labels, this leads to a complicated algorithm with an exponential time complexity. However, we have a linear-time algorithm for the use case that is most important from the perspective of this study: comparing species and gene trees.

This article focuses on the algorithmical aspects of the Branch Displacement Index and contains two examples that serve as an illustration of its application and properites. Further and more detailed studies on how its performance compares to other methods, and on its theoretical properties in different evolutionary scenarios, would be desirable. For example, a rooted tree is much more informative than an unrooted one, so naturally, we lose some capabilities when we only use unrooted trees. In particular, some horizontal gene transfers are undetectable in this setting. For example, in Fig. 1, a transfer from A to X would not change the branching patterns in the trees, but it would be detectable in a rooted gene tree because the X gene would branch with A instead of being an outgroup. Whether this drawback is outweighted by the potentially reduced false positive rate is yet to be elucidated.

Another application of CDI is to evaluate the consistency of the location of a given clade in bootstrap replicates. Bootstrap support values (and other similar measures) give us infomation about the existence of a given clade; it is, however, far from straightforward to use them to determine whether the clade’s location is supported by the data. This is most clearly visible in the case of leaves: any given leaf always exists as a clade, but its location may vary greatly between bootstrap replicates. In general, an unstable location of a clade lowers the bootstrap values of its incident edges, not the edge that defines the clade.

On a final note we point that, unlike the gene trees, the species trees are typically rooted. Our approach, however, does not use the information about the root location of the species tree, which may potentially be used to improve the specificity and/or sensitivity of our approach. The reason for not using this information is simple: as for now, we have not come up with an idea how.

## Acknowledgements

We thank Paweł Górecki and Anna Gambin from the faculty of Mathematics, Informatics and Mechanics, University of Warsaw and Anna Muszewska from the Institute of Biochemistry and Biophysics, Polish Academy of Sciences for reading the draft of this manuscript and thoughtful remarks.

The support for this work was provided by the Polish National Science Centre grant 2019/33/B/ST6/00737.

## Notes

https://github.com/mciach/CDI

### Competing Interest Statement

The authors have declared no competing interest.

### Summary of Updates

Title of the manuscript updated to eliminate a typo

## References

[1] Matt Ravenhall, Nives Škunca, Florent Lassalle, and Christophe Dessimoz. Inferring horizontal gene transfer. PLoS Comput Biol, 11(5):e1004095, 2015.

[2] Shannon M Soucy, Jinling Huang, and Johann Peter Gogarten. Horizontal gene transfer: building the web of life. Nature Reviews Genetics, 16(8):472–482, 2015.

[3] Jennifer Becq, Cécile Churlaud, and Patrick Deschavanne. A benchmark of parametric methods for horizontal transfers detection. PLoS One, 5(4):e9989, 2010.

[4] Corinne Rancurel, Ludovic Legrand, and Etienne GJ Danchin. Alienness: rapid detection of candidate horizontal gene transfers across the tree of life. Genes, 8(10):248, 2017.

[5] Patrick J Keeling and Jeffrey D Palmer. Horizontal gene transfer in eukaryotic evolution. Nature Reviews Genetics, 9(8):605–618, 2008.

[6] Maria S Poptsova and J Peter Gogarten. The power of phylogenetic approaches to detect horizontally transferred genes. BMC evolutionary biology, 7(1):1–17, 2007.

[7] Nancy A Moran and Tyler Jarvik. Lateral transfer of genes from fungi underlies carotenoid production in aphids. science, 328(5978):624–627, 2010.

[8] Satoko Yoshida, Shinichiro Maruyama, Hisayoshi Nozaki, and Ken Shirasu. Horizontal gene transfer by the parasitic plant striga hermonthica. Science, 328(5982):1128–1128, 2010.

[9] Nora Khaldi, Jérôme Collemare, Marc-Henri Lebrun, and Kenneth H Wolfe. Evidence for horizontal transfer of a secondary metabolite gene cluster between fungi. Genome biology, 9(1):1–10, 2008.

[10] Boran Altincicek, Jennifer L Kovacs, and Nicole M Gerardo. Horizontally transferred fungal carotenoid genes in the two-spotted spider mite tetranychus urticae. Biology letters, 8(2):253–257, 2012.

[11] L Yu Rusin, EV Lyubetskaya, K Yu Gorbunov, and VA Lyubetsky. Reconciliation of gene and species trees. BioMed research international, 2014, 2014.

[12] Jean-Philippe Doyon, Vincent Ranwez, Vincent Daubin, and Vincent Berry. Models, algorithms and programs for phylogeny reconciliation. Briefings in bioinformatics, 12(5):392–400, 2011.

[13] Maureen Stolzer, Han Lai, Minli Xu, Deepa Sathaye, Benjamin Vernot, and Dannie Durand. Inferring duplications, losses, transfers and incomplete lineage sorting with nonbinary species trees. Bioinformatics, 28(18):i409–i415, 2012.

[14] Yi-Chieh Wu, Matthew D Rasmussen, Mukul S Bansal, and Manolis Kellis. Most parsimonious reconciliation in the presence of gene duplication, loss, and deep coalescence using labeled coalescent trees. Genome research, 24(3):475–486, 2014.

[15] Lars Arvestad, Ann-Charlotte Berglund, Jens Lagergren, and Bengt Sennblad. Bayesian gene/species tree reconciliation and orthology analysis using mcmc. Bioinformatics, 19(suppl 1):i7–i15, 2003.

[16] Hyeonsoo Jeong, Samsun Sung, Taehyung Kwon, Minseok Seo, Kelsey Caetano-Anollés, Sang Ho Choi, Seoae Cho, Arshan Nasir, and Heebal Kim. Hgtree: database of horizontally transferred genes determined by tree reconciliation. Nucleic acids research, 44(D1):D610–D619, 2016.

[17] Lawrence A David and Eric J Alm. Rapid evolutionary innovation during an archaean genetic expansion. Nature, 469(7328):93–96, 2011.

[18] Pawel Górecki, Oliver Eulenstein, and Jerzy Tiuryn. Unrooted tree reconciliation: A unified approach. IEEE/ACM transactions on computational biology and bioinformatics, 10(2):522–536, 2013.

[19] Agnieszka Mykowiecka and Paweł Górecki. Credibility of evolutionary events in gene trees. IEEE/ACM transactions on computational biology and bioinformatics, 16(3):713–726, 2018.

[20] Ziheng Yang and Bruce Rannala. Molecular phylogenetics: principles and practice. Nature reviews genetics, 13(5):303–314, 2012.

[21] Stéphane Guindon, Jean-François Dufayard, Vincent Lefort, Maria Anisimova, Wim Hordijk, and Olivier Gascuel. New algorithms and methods to estimate maximum-likelihood phylogenies: assessing the performance of phyml 3.0. Systematic biology, 59(3):307–321, 2010.

[22] Bui Quang Minh, Heiko A Schmidt, Olga Chernomor, Dominik Schrempf, Michael D Woodhams, Arndt Von Haeseler, and Robert Lanfear. Iq-tree 2: New models and efficient methods for phylogenetic inference in the genomic era. Molecular biology and evolution, 37(5):1530–1534, 2020.

[23] Tom A Williams, Sarah E Heaps, Svetlana Cherlin, Tom MW Nye, Richard J Boys, and T Martin Embley. New substitution models for rooting phylogenetic trees. Philosophical Transactions of the Royal Society B: Biological Sciences, 370(1678):20140336, 2015.

[24] Mathieu Groussin, Bastien Boussau, and Manolo Gouy. A branch-heterogeneous model of protein evolution for efficient inference of ancestral sequences. Systematic biology, 62(4):523–538, 2013.

[25] Seraina Klopfstein, Lars Vilhelmsen, and Fredrik Ronquist. A nonstationary markov model detects directional evolution in hymenopteran morphology. Systematic biology, 64(6):1089–1103, 2015.

[26] Tonny Kinene, J Wainaina, Solomon Maina, and LM Boykin. Rooting trees, methods for. Encyclopedia of Evolutionary Biology, page 489, 2016.

[27] Uyen Mai, Erfan Sayyari, and Siavash Mirarab. Minimum variance rooting of phylogenetic trees and implications for species tree reconstruction. PloS one, 12(8):e0182238, 2017.

[28] Fernando Domingues Kümmel Tria, Giddy Landan, and Tal Dagan. Phylogenetic rooting using minimal ancestor deviation. Nature Ecology & Evolution, 1(1):1–7, 2017.

[29] Ran Libeskind-Hadas, Yi-Chieh Wu, Mukul S Bansal, and Manolis Kellis. Pareto-optimal phylogenetic tree reconciliation. Bioinformatics, 30(12):i87–i95, 2014.

[30] Taylor Wade, L Thiberio Rangel, Soumya Kundu, Gregory P Fournier, and Mukul S Bansal. Assessing the accuracy of phylogenetic rooting methods on prokaryotic gene families. PloS one, 15(5):e0232950, 2020.

[31] Soumya Kundu and Mukul S. Bansal. On the impact of uncertain gene tree rooting on duplication-transfer-loss reconciliation. BMC Bioinformatics, 19(S9), August 2018.

[32] Alessandra P Lamarca and Carlos G Schrago. Fast speciations and slow genes: uncovering the root of living canids. Biological Journal of the Linnean Society, 129(2):492–504, 2020.

[33] Laura M Boykin, Laura Salter Kubatko, and Timothy K Lowrey. Comparison of methods for rooting phylogenetic trees: A case study using orcuttieae (poaceae: Chloridoideae). Molecular phylogenetics and evolution, 54(3):687–700, 2010.

[34] David F Robinson and Leslie R Foulds. Comparison of phylogenetic trees. Mathematical biosciences, 53(1-2):131–147, 1981.

[35] Sebastian Böcker, Stefan Canzar, and Gunnar W Klau. The generalized robinson-foulds metric. In International Workshop on Algorithms in Bioinformatics, pages 156–169. Springer, 2013.

[36] Robert J Vanderbei et al. Linear programming, volume 3. Springer, 2015.

[37] Kazutaka Katoh and Daron M Standley. Mafft multiple sequence alignment software version 7: improvements in performance and usability. Molecular biology and evolution, 30(4):772–780, 2013.

[38] Jose Castresana. Selection of conserved blocks from multiple alignments for their use in phylogenetic analysis. Molecular biology and evolution, 17(4):540–552, 2000.

